# NFAT5-dependent transcriptional stress control of endothelial cells prevents maladaptive remodeling of pulmonary arterioles in the hypoxic lung

**DOI:** 10.1101/2023.10.18.563022

**Authors:** Hebatullah Laban, Sophia Sigmund, Katharina Schlereth, Lennart Brandenburg, Felix A. Trogisch, Andreas Weigert, Carolina De La Torre, Alia Abukiwan, Carolin Mogler, Markus Hecker, Wolfgang M. Kuebler, Thomas Korff

**Affiliations:** Institute of Physiology and Pathophysiology, Department of Cardiovascular Physiology, Heidelberg University, Germany; Deutsches Zentrum für Herz-Kreislauf-Forschung e.V. (DZHK), Partner Site Heidelberg/Mannheim, 69120 Heidelberg, Germany; Division of Vascular Oncology and Metastasis, German Cancer Research Center (DKFZ-ZMBH Alliance), Heidelberg, Germany; European Center for Angioscience (ECAS), Medical Faculty Mannheim, Heidelberg University, Germany; Department of Cardiovascular Physiology and Cardiac Imaging Center, Core Facility Platform Mannheim, Medical Faculty Mannheim, Heidelberg University, Germany; Institute of Biochemistry I Pathobiochemistry, Faculty of Medicine, Goethe University, Frankfurt am Main, Germany; NGS Core Facility, Medical Faculty Mannheim, Heidelberg University, Germany; Institute of Pathology, School of Medicine, Technical University Munich, Germany; Institute of Physiology, Charité-Universitätsmedizin Berlin, corporate member of Freie Universität Berlin and Humboldt Universität zu Berlin.

## Abstract

**Aims:** Chronic hypoxia causes detrimental structural alterations in the lung, which are partially dependent on stress responses of the endothelium. In this context, we revealed that hypoxia-exposed murine lung endothelial cells (MLEC) activate nuclear factor of activated T-cells 5 (NFAT5) - a transcription factor that adjusts the cellular transcriptome to cope with multiple environmental stressors. Here, we studied the functional relevance of NFAT5 for the control of hypoxia-induced transcription in MLEC.

**Methods and Results:** Targeted ablation of *Nfat5* in endothelial cells did not evoke phenotypic abnormalities in normoxia-exposed mice. However, MLEC in *Nfat5*-deficient mice up-regulated energy- and protein-metabolism-associated gene expression under normobaric hypoxia (10% O_2_) for seven days as evidenced by microarray- and scRNA-seq-based analyses. Moreover, loss of NFAT5 boosted the expression and release of platelet-derived growth factor B (*Pdgfb)* - a HIF1α-regulated driver of vascular smooth muscle cell (VSMC) growth - in capillary MLEC of hypoxia-exposed mice, which was accompanied by exaggerated coverage of distal pulmonary arterioles by VSMC, increased pulmonary vascular resistance and impaired right ventricular functions. *In vitro,* knockout of *Nfat5* in cultured MLEC stimulated *Pdgfb* expression and release after exposure to hypoxia and amplified binding of HIF1α in the *Pdgfb* promoter region.

**Conclusion:** Collectively, our study identifies NFAT5 as a protective transcription factor required to rapidly adjust the transcriptome of MLEC to hypoxia. Specifically, NFAT5 restricts HIF1α-mediated *Pdgfb* expression and consequently limits muscularization and resistance of pulmonary arterioles.

**Highlights:**

- Hypoxia stimulates the transcriptional activity of NFAT5 in MLEC.
- Loss of NFAT5 in hypoxia-exposed MLEC results in EC subtype-specific maladaption of growth factor-, energy- and protein-metabolism-associated gene expression.
- Specifically, NFAT5-deficient capillary lung EC unleash HIF1α-regulated *Pdgfb* expression and release, which results in excessive coverage of pulmonary arterioles by VSMC.
- NFAT5-dependent control of early stress responses of capillary MLEC is required to limit the increase in pulmonary vascular resistance and impairment of right ventricular functions.

**Graphical Abstract:** 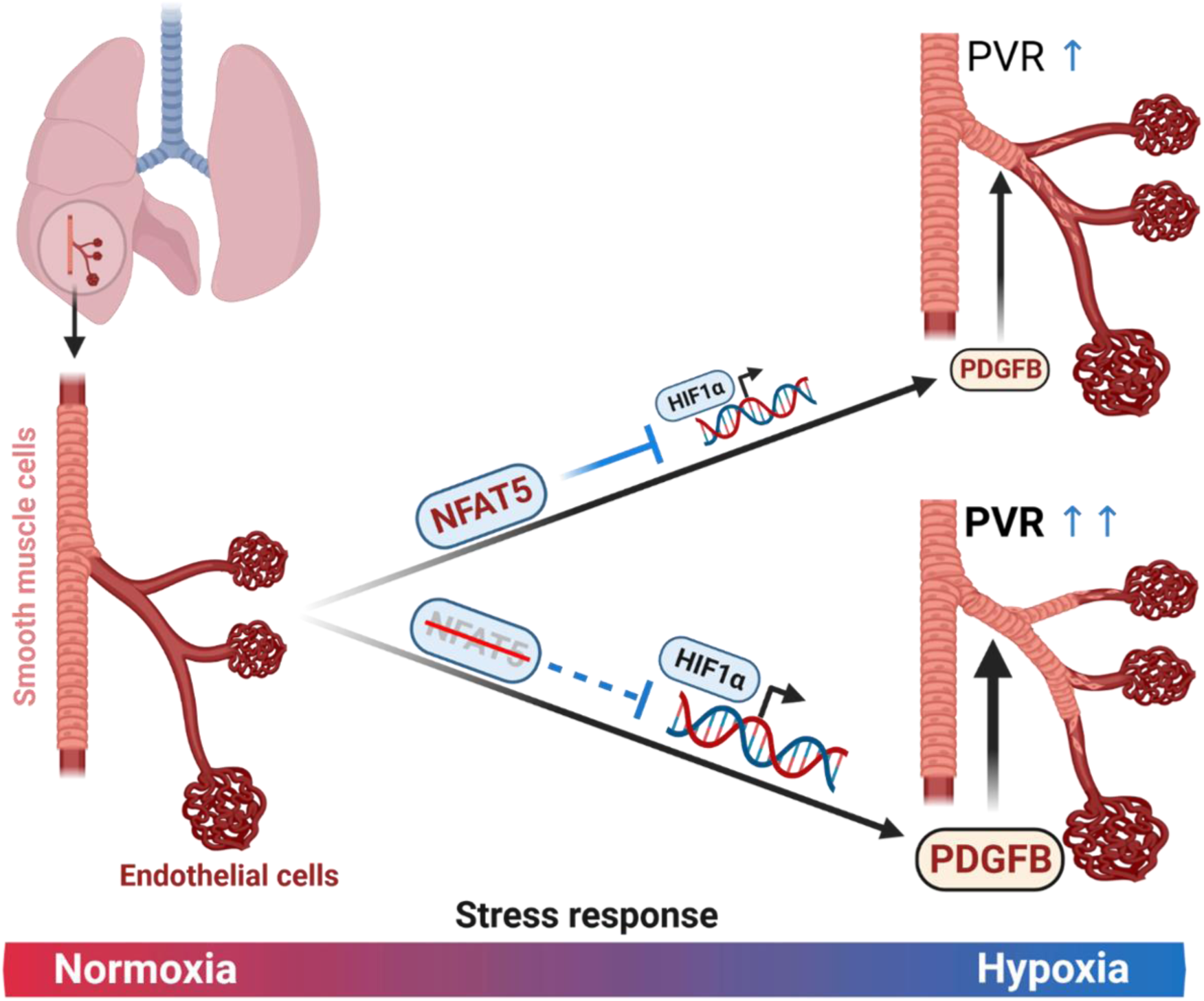

## INTRODUCTION

The mammalian organism has several options to deal with hypoxia. For instance, oxygen delivery may be improved by increasing heart rate and erythropoiesis. In addition to such systemically acting mechanisms, organ-specific adaptive reactions are triggered. One of the foremost tissues responding to a hypoxic environment are pulmonary arteri(ol)es and capillaries, which transport deoxygenated blood from the right ventricle to the alveoli. In fact, a drop in oxygen partial pressure (pO_2_) promotes constriction of small pulmonary arteri(ol)es to redirect blood flow to other areas of the lung for maintained blood oxygenation. This response is known as hypoxic pulmonary vasoconstriction (HPV)^1^ and matches capillary perfusion to alveolar ventilation ^2^. However, chronic exposure to hypoxic conditions as evoked by chronic obstructive pulmonary disease (COPD) or hypoventilation triggers additional and long-lasting structural changes of pulmonary blood vessels including loss of precapillary arteries, medial hypertrophy of large pulmonary arteries and (ectopic) coverage of normally not muscularized distal arterioles by VSMCs ^3, 4^. The latter abnormality is one of the most common finding in mouse models of hypoxia ^5^. In this context, endothelial cells (EC) appear to initiate the activation of VSMC by expressing platelet-derived growth factor subunit B (*Pdgfb)* under the control of hypoxia-inducible factor 1 alpha (HIF1A). PDGFB stimulates the activity of the Krüppel-like (transcription) factor 4 (KLF4) in VSMCs as a crucial determinant of their growth and migration ^6^. This process was observed seven days after the onset of hypoxia and originated from a specific VSMC progenitor population that acquires an activated phenotype to migrate and proliferate, finally returning to a quiescent differentiated state ^7, 8^.

Both, the (initial) HPV and the degree of VSMC coverage progressively increase the resistance of (distal) pulmonary arteri(ol)es and consequently the afterload for the right ventricle. Patients affected by these pathophysiological events usually suffer from pulmonary arterial hypertension (PAH) accompanied by right ventricular hypertrophy and ultimately right heart failure. Despite the discovery of several fundamental steps of the pathogenesis, therapeutic options to face PAH are still limited. Thus, understanding of molecular mechanisms increasing the resistance of pulmonary arterial system appears of utmost relevance for developing effective treatment strategies in the future.

We recently reported that hypoxia stimulates the transcriptional activity of nuclear factor of activated T-cells 5 (NFAT5) in pulmonary artery VSMC of the mouse lung^9^. Under these conditions, NFAT5 regulated the cellular transcriptome to cope with hypoxia, thereby limiting pulmonary vascular resistance by supporting hypoxia-adapted mitochondrial function ^9^. NFAT5 or tonicity enhancer binding protein (TonEBP) is a calcineurin-independent member of the Rel family of transcription factors and has been shown to regulate multiple types of cellular stress responses associated with hyperlipidemia, insulin resistance, arteriosclerosis, rheumatoid arthritis, hypertonicity ^10, 11^ and ischemia ^12^. In vascular EC, NFAT5 was reported to promote their angiogenic and pro-inflammatory activation ^13^ but its impact on the hypoxia-induced endothelial stress response in the lung is unknown. Considering the functional relevance of NFAT5 for pulmonary VSMC in the hypoxic lung, we hypothesized that it may similarly affect the transcriptome of pulmonary EC, enabling them to adequately cope with a hypoxic environment. Consequently, we generated mice with an EC-specific ablation of *Nfat5*, exposed them to normobaric hypoxia and investigated the gene expression profile of MLEC *in vivo* and *in vitro*.

## MATERIAL AND METHODS

### Cell culture

Murine lung endothelial cells (MLEC) were isolated from adult Nfat5^fl/fl^ mice as the follows: Lungs were dissected, and the lobes were carefully minced with a scalpel and digested with 4 U/ml dispase II in Ham’s F12 for 1 h at 37°C. The digested tissue was homogenized by pipetting and filtered through a 70 µm cell strainer into Ham’s F12 containing 10% fetal calf serum (FCS). After washing, CD144-positive endothelial cells were sorted using rat anti-mouse CD144 antibody (BD Biosciences, #555289, Heidelberg, Germany) -coated magnetic beads (Dynabeads, Invitrogen, #11035, Waltham, Massachusetts, USA). Isolated CD144^+^ MLEC were grown on fibronectin-coated culture plates and cultured in DMEM/F12 medium supplemented with 20% FCS, 4–8 µl/ml ECGS-H, 50 U/ml penicillin, and 50 µg/ml streptomycin at 37°C in an atmosphere with 5% CO_2_. MLEC were purified after propagating them two times by sorting CD31^+^/CD144^+^ cells as described above (anti-mouse CD31-antibody, BD Biosciences, #553370, Heidelberg, Germany). Cell identity and purity were routinely checked by performing immunofluorescence-based or FACS analysis to detect endothelial cell marker CD31 (Supplement 1). For some experiments, MLEC were pretreated with 30 µM of DNA methyltransferase inhibitor (RG108, sc-204235, Santa Cruz, Biotechnology, Heidelberg, Germany) or 10 µM HIF-1α inhibitor (CAY10585, Cat# sc-205346, Santa Cruz Biotechnology, Heidelberg, Germany) for 1 h. Treatment with equal concentration of solvent served as control. MLEC were exposed to hypoxia (1% O_2_) for up to 24 h in a hypoxia chamber (Invivo2 Plus, Ruskinn).

### Viral transduction

Deletion of the floxed *Nfat5* exon was achieved by transduction with adenoviral vectors to express Cre recombinase (iCre) under the control of a cytomegalovirus (CMV) promoter (Ad-CMV-iCre, Cat. # 1045, Vector Biolabs, Malvern, PA, USA). The virus was applied to the cellswith a multiplicity of infection (MOI) of 500. An adenovirus with an empty CMV promoter (Ad-CMV-Null, Cat. # 1300, Vector Biolabs, Malvern, PA, USA) was used as a control. Cells were used for experiments 48 hours after exposure to adenoviral vectors. Knockout efficiency was monitored by determining the *Nfat5* expression and protein levels (Supplement 2).

Overexpression of *Nfat5* in MLEC was achieved by transducing them with an adenoviral vector expressing human NFAT5c under a CMV promoter (Sirion Biotech, Munich, Germany) with a MOI of up to 100. A control plasmid was purchased from Sirion Biotech (Germany). Human *NFAT5* shows 92% protein sequence homology as compared to murine *Nfat5* and 99% similarity for the NH2-terminal 500 amino acids, which include the Rel-like DNA binding domain ^14^.

### Immunofluorescence analyses of cultured cells

MLEC were rinsed with HBSS and fixed with ice-cold methanol for 15 minutes and allowed to dry for 20 minutes. They were treated with 0.5 % Triton-X and 10% donkey serum in PBS for 30 minutes, and incubated with rabbit anti-NFAT5 antibody 1:100 (Invitrogen, PAI-023, Waltham, Massachusetts, USA) with or without goat anti-CD31 (Biotechne, AF3628, Wiesbaden-Nordenstadt, Germany) at 4°C overnight. After washing, MLEC were incubated with appropriate fluorophore-labelled secondary antibodies for 1 hour and mounted with Mowiol (Calbiochem, San Diego, CA, USA). Nuclei were visualized by DAPI staining. Fluorescence images were acquired by an AxioVert 200M microscope (Zeiss) using the TissueFAXS software (version 4.2) and TissueQuest software (version 4.0.1.0137) for automated image analysis (TissueGnostics GmbH, Vienna) and on an Olympus IX83 microscope (Olympus, Hamburg, Germany) using cellSens software (v. 1.12).

### Analysis of gene expression

RNA was extracted from MLEC and lung tissue by lysis in RLT buffer containing 1% β-mercaptoethanol for RNA isolation and subsequently purified using the RNeasy Mini Kit (Qiagen, 74106, Hilden, Germany) according to manufacturer’s instructions. RNA from FAC-sorted lung EC was extracted using the Nucleaospin^®^ TriPrep Kit (MACHEREY-NAGEL, 740966.50, Düren, Germany Germany). RNA quantity and quality were assessed using the NanoDrop ND-1000 spectrophotometer (NanoDrop Technologies, Wilmington, DE, USA). cDNA was synthesized using the Sensiscript Reverse Transcription Kit (Qiagen, 205213, Hilden, Germany) or the Omniscript Reverse Transcription Kit (Qiagen, 205113, Hilden, Germany) and oligo(dT)15 primers (Promega, C110A, Madison, WI, USA). Quantitative real-time RT-PCR was performed in a Rotor-Gene Q (Qiagen, Hilden, Germany) using the 5x QPCR Mix EvaGreen (No Rox) (Solis Biodyne, Cat. # 08-25-00001-20, Estonia) and purified water (Merck Millipore, Cat# H20MB0106, Darmstadt, Germany). PCR was conducted in 40 cycles of denaturation (95°C, 15 s), annealing (AT, temperature depending on the primer, 25 s) and elongation (72°C, 10 s) after one initial phase of 95°C for 15 min to activate the DNA polymerase. Fluorescence was assessed at the end of the annealing phase (excitation at 470 nm, emission at 530 nm). The threshold cycle (Ct) was set within the exponential phase of the PCR using the Rotor-Gene Q Series software (v.2.3.1, Build 49, Qiagen, Hilden, Germany). The PCR product was quantified using the 2^−ΔΔCt^ method. Amplification of *Actb* was used as an internal standard. QuantiTecht Primer Assays (Qiagen, Germany) were utilized for *Pdgfb* (QT00266910, #249900) and *Tgfb1* (QT00145250, #307447906). The following primers were used: *Nfat5* forward 5’-GGGGGTCAAACGACGAGATT-3’ and reverse 5’-ACTGTCCGCACAACATAGGG-3’,(annealing temperature (AT: 59 °C). *Actb* forward 5’-CGGTTCCGATGCCCTGAGGCTCTT-3’ and reverse 5’-CGTCACACTTCATGATGGAATTGA-3’ (AT: 60 °C) and *Vegfa* forward 5’-CAGGCTGCTGTAACGATGAA-3’ and reverse 5’-GCATTCACATCTGCTGTGCT-3’, (AT: 60 °C), *Rps12* forward 5’-GAAGCTGCCAAAGCCTTAGA-3’ and reverse 5’-AACTGCAACCAACCACCTTC-3’ (AT: 60 °C), *Cxcl12* forward 5’-GGAGGATAGATGTGCTCTGGAAC-3’ and reverse 5’-AGTGAGGATGGAGACCGTGGTG-3’ (AT: 60 °C).

### Genome array-based analyses

Gene expression profiling of RNA isolated from whole lung tissue was performed using the GeneChip® Mouse Gene 2.0 ST Array (902118, Affymetrix). A custom CDF version 22 with ENTREZ-based gene definitions was used to annotate the arrays. Raw fluorescence intensity values were normalized using quantile normalization. Pathways associated with different cell functions were obtained from external public databases (KEGG, http://www.genome.jp/kegg). The raw microarray data are available in the Gene Expression Omnibus (GEO) database as record GSE230055 at https://www.ncbi.nlm.nih.gov/geo/query/acc.cgi?acc=GSE230055, and can be accessed for review using the following passcode: evanwmcwrzwfxmv

### Capillary electrophoresis and protein analyses

Whole cell lysates from MLEC or from frozen lung lobes were prepared in radioimmunoprecipitation assay (RIPA) buffer. Lysates of cytosolic or nuclear fractions were prepared by lysing cells in buffer I containing 10 mM HEPES, 10 mM KCl, 1 µM EDTA, 1 µM EGTA, 15 % nonidet, protease and phosphatase inhibitors. After centrifugation (12,000 x g at 4 °C for 15 min) the supernatant containing the cytosolic protein fraction was used for further analyses. The remaining sediment was dissolved in 40 µl of buffer II consisting of 20 mM HEPES, 400 mM NaCl, 0.01 M EDTA, 0.01 M EGTA, 15% nonidet and protease and phosphatase inhibitors. Subsequently, this solution was exposed to ultrasound (50 W) twice for 5 seconds at 4°C. After centrifugation (12,000 x g at 4°C for 15 minutes), the supernatant containing the nuclear protein fraction was used for further analyses.

Lysates were processed for capillary electrophoresis using Simple Western technology (WES, ProteinSimple, Bio-Techne, San Jose, CA, USA). Sample preparation and antibody application were performed according to the manufacturer’s instructions using the 12 - 230 kDa WES separation module (Cat. #. SM-W004, ProteinSimple, Bio-Techne). The following antibodies were used (Cat. # supplier, dilution) as the following: anti-NFAT5 (sc-398171, Santa Cruz, Biotechnology, Heidelberg, Germany 1:10), anti-α-tubulin (CS#2144, Cell Signaling, 1:10), anti-HDAC1 (NB100-56340, Novus Biologicals, 1:10), anti-Hif1α (CS#14179s, Cell Signaling, Danvers, MA, USA, 1:150) and anti-β-actin (MAB8929, R&D Systems, Abingdon, UK, 1:500). β-actin served as loading reference. Protein levels were determined using the Compass software (version 4.0.1, build 0812, ProteinSimple, Bio-Techne, San Jose, CA, USA).

### Chromatin immunoprecipitation (ChIP) assay

MLEC were exposed to hypoxia for 8 hours, detached, suspended and fixed in formaldehyde (final conc. 1% in PBS) on a rocking platform to fix the protein-DNA interactions. After 10 min, 1.25 M Glycine was added and the cross-linked cells was lysed to release the chromatin containing DNA-protein complexes. Chromatin was isolated using a ChIP Kit (Cat. #ab185913, Abcam, Trumpington, UK). Chromatin was fragmented into smaller pieces by ultrasound according to manufacturer’s instructions. Immunoprecipitation was performed with anti-NFAT5 (sc-398171, Santa Cruz, Biotechnology, Heidelberg, Germany), or anti-Hif1α (CS#14179s, Cell Signaling, Danvers, MA, USA) or non-immune IgG. One µg of the antibodies was allowed to bind to the wells of test strip for 90 min at RT. After washing, 2 µg chromatin solution was added in each well and incubated overnight at 4°C. Reversal of cross-linking and DNA purification were performed according to manufacturer’s instructions. Quantity and quality of the purified DNA were assessed using the NanoDrop ND-1000 spectrophotometer (NanoDrop Technologies, Wilmington, DE, USA). Five ng of purified DNA was used for qRT-PCR analysis in a Rotor-Gene Q (Qiagen, Hilden, Germany) using the 5x QPCR Mix EvaGreen (No Rox) (Solis Biodyne, Cat. # 08-25-00001-20, Estonia) and purified water (Merck Millipore, Cat# H20MB0106, Darmstadt, Germany). PCR was conducted in 40 cycles including i) denaturation (95°C, 15 s), ii) annealing (AT, temperature depending on the primer, 25 s) and iii) elongation (72°C, 10 s) after one initial phase of 95°C for 15 min. Fold enrichment (FE) was calculated as the ratio of amplification efficiency of the ChIP sample to non-immune IgG (FE% = 2 ^(IgG^ ^CT-Sample^ ^CT)^ X 100%). The sequence of a NFAT5 binding site (NP-4) in the *Pdgfb* promoter region that overlaps with a hypoxia response element (HRE) was identified by applying the Genomatrix software suite v3.13 (MatInspector, Precigen Bioinformatics, Munich, Germany; Supplement 3). Sequences of NP-4-specific primers were: NP-4 forward 5’-CCACTTAGAGAATGTAGG-3’ and reverse 5’-GCTTAGATTCCATTGAGT - 3’ (AT: 60 °C).

### Methylation assay

DNA from MLEC was extracted, purified and analyzed to quantify 5-methylcytosine levels by applying the DNA Purification Kit (T3010S, Monarch, New England Biolabs, USA) and the 5mC Assay Kit (lot no. ab117128, Abcam, Trumpington, UK) following manufacturer’s instructions. Global DNA methylation was determined by colorimetric measurements using a BioTek Absorbance Reader.

### Assay for Transposase-Accessible Chromatin using sequencing (ATAC-seq)

MLEC were detached, suspended and counted. Equal numbers of cells per sample were frozen in liquid nitrogen and sent to Active Motif (USA) for further ATAC-seq analyses. Initial adapter sequence trimming of the FASTQ files was followed by mapping to the genome using the BWA algorithm. Only high-quality, uniquely aligned reads were retained for subsequent analysis, with removal of duplicate reads. Peaks were identified using the MACS2 algorithm, applying a P-value threshold of < 10^-10^ for filtering. Peak raw counts were quantified, and intervals were defined based on chromosome number and start/end coordinates. Overlapping intervals were grouped into “merged regions” to enable comparisons of peak metrics between samples. Merged regions were determined by the start and end coordinates of the overlapping intervals, facilitating fragment density calculations even in samples without called peaks. Data visualization was performed using the Integrated Genome Browser ^15^.

### Mouse model

All animal studies were performed with the approval of the Regional Council Karlsruhe (approval number: 35-9185.81/G-233/18) and in accordance with the Guide for the Care and Use of Laboratory Animals published by the US National Institutes of Health (NIH Publication No. 85-23, revised 1996) as well as Directive 2010/63/EU of the European Parliament on the protection of animals used for scientific purposes. All mice were housed according to GV-SOLAS recommendations and were hygienically monitored by sentinel animals. *Nfat5^fl/fl^* mice were described previously ^16^ and crossed with *Cdh5*-CreERT2 mice ^17^. Genetic ablation (deletion of loxP site-flanked exon 4 of *Nfat5*) of *Nfat5* (*Nfat5^(EC)-/-^*) in the corresponding offspring was induced at 10-12 weeks of age by i.p. administration of 1 mg tamoxifen per day for 5 consecutive days or miglyol as solvent control (*Nfat5^fl/fl^*). Female and male Mice were used in experiments after a recovery period of 2-3 weeks.

Mice were exposed to normobaric hypoxia for 7 or 21 days by placing them in a hypoxia chamber (A-Chamber/ProOx O_2_ controller, Biospherix, USA) with free access to drinking water and food (humidity: 50-65%, temperature: 21-23°C) with 10% oxygen and 90% nitrogen. To detect proliferating cells, mice were treated with a single dose of 2 mg 5-ethynyl-2′-deoxyuridine (EdU, 900584, Sigma Aldrich) in PBS by intraperitoneal injection 2 h before euthanasia by cervical dislocation. Mice showing excessive weight loss or pathologic lethargy after tamoxifen treatment or while being exposed to hypoxia were excluded from further analyses.

### Isolation of MLEC by FACS

Isolation of MLEC was performed as based on a method previously described ^18^. Briefly, lungs were digested by applying the Lung Dissociation Kit (Miltenyi Biotec, Bergisch Gladbach, Germany) following manufacturer’s instruction. Single-cell suspensions were prepared by passing the digestion mixture through 18G and 19G cannula syringes and filtered through a 100 µm cell strainer. Erythrocytes were lysed after the addition of ACK buffer (154.4 mM ammonium chloride, 10 mM potassium bicarbonate, and 97.3 μM EDTA tetrasodium salt). CD31^high^ cells were enriched by magnetic activated cell sorting (MACS). To exclude non-(vascular) endothelial cell populations, cell suspensions were treated with the following antibodies: anti-PDPN-AlexaFluor® 488 (eBioscience, Frankfurt am Main, Germany, 53–5381, RRID: AB_1106990), anti-LYVE1-AlexaFluor® 488 (eBioscience, 53– 0443, RRID:AB_1633415), anti-CD45-FITC (BD Biosciences, Heidelberg, Germany, 553080, RRID:AB_394609), anti-LY76-FITC (BD Biosciences, 561032, RRID:AB_396936) and anti-EPCAM-FITC (eBioscience, 11-5791, RRID:AB_ 11151709) for 30 min at 4°C in PBS containing 5% FCS. Vascular endothelial cells were simultaneously labelled with anti-CD31-APC (BD Biosciences, 551262, RRID: AB_398497) and anti-CD146-PE-Cy7 (BioLegend, San Diego, CA, USA, 134713, RRID: AB_2563108). Dead cells were excluded by propidium iodide (PI, 1:2000) staining. CD45^-^ LYVE1^-^ LY76^-^ PDPN^-^ EPCAM^-^ PI^-^ CD31^+^ CD146^+^ cells were sorted using a BD FACSAria Fusion Flow Cytometer (BD Biosciences, Heidelberg, Germany). Immediately after sorting, cells were frozen in DMSO-supplemented MLEC medium on dry ice or directly processed for further analyses.

### Flow cytometry-based analyses

Single-cell suspensions were prepared from mouse lungs using the Lung Dissociation Kit (Miltenyi Biotec, Bergisch Gladbach, Germany) following manufacturer’s instructions. Cells were incubated with anti-CD3e (BD Biosciences, Heidelberg, Germany, 557757, RRID:AB_396863), anti-CD4 (BD Biosciences, 560782, RRID:AB_1937315 and 563050, RRID:AB_2737973), anti-CD8a (Biolegend, San Diego, CA, USA, 100742, RRID:AB_2563056), anti-CD11b (Biolegend, 101257, RRID:AB_2565431), anti-CD11c (BD Biosciences, 563048, RRID:AB_2734778), anti-CD24 (BD Biosciences, 562477, RRID: AB_11151917), anti-CD44 (BD Biosciences, 560567, RRID:AB_1727480), anti-CD45 (Miltenyi, 130-102-430, RRID: AB_2659925), anti-SiglecF (BD Biosciences, 562068, RRID:AB_10896143), anti-MerTK (Biolegend, 151521, RRID:AB_2876508), anti-HLA-DR (MHCII) (Miltenyi Biotec, Bergisch Gladbach, Germany, 130-102-139, RRID: AB_2660058), anti-Ly6C (BD Biosciences, 560525, RRID:AB_1727558), anti-Ly6G (Biolegend, 127624, RRID:AB_10640819), anti-SiglecH (Biolegend, 129604, RRID: AB_1227761), anti-CD31 (eBioscience, Frankfurt am Main, Germany, 25-0311-82, RRID:AB_2716949), anti-CD90.2 (Miltenyi, 130-102-960, RRID:AB_2659874), anti-CD140b (Biolegend, 136006, RRID:AB_1953271), anti-CD146 (Biolegend, 134708, RRID:AB_10640123), anti-CD324 (Biolegend, 147308, RRID:AB_2563955), anti-CD326 (BD Biosciences, 563134, RRID:AB_2738022), anti-GITR (CD357) (Biolegend, 126308, RRID:AB_1089125), anti-NK1.1 (BD Biosciences, 563096, RRID:AB_2738002) and anti-TCR γδ (Biolegend, 118116, RRID:AB_1731813). Antibodies and secondary reagents were titrated to determine optimal concentrations. CompBeads (BD Biosciences) were used for single-color compensation to create multicolor compensation matrices. Fluorescence minus one (FMO) controls were used for gating. The instrument calibration was controlled daily using Cytometer Setup and Tracking beads (BD Biosciences, Heidelberg, Germany). Samples were analyzed on a LSRII/Fortessa flow cytometer (BD Biosciences, Heidelberg, Germany) and the data were processed using FlowJo (V10.6.1., Treestar; see Supplement 4 for gating strategy)

### 10x Genomics Transcriptomics (scRNAseq)

Lungs were digested in DMEM/F12 (ThermoFisher Scientific, Germany) containing 4 U/ml Dispase (Sigma, Munich, Germany) and 10 µg/ml DNaseI (Roche, Germany) at 37°C for 45 min. Digested tissues were filtered through a 100 µm cell strainer. Erythrocytes were lysed using ACK buffer. Anti-mouse CD16/32 dimer antibody was used to inhibit non-specific binding of antibodies to FcγII and FcγIII. The cell suspension was incubated with anti-CD31 MicroBeads (Miltenyi Biotec, Germany) to pre-select PECAM-1^high^ cells using MACS LS columns and a magnetic separator. Subsequently, CD45^low^ LYVE1^low^ LY76^low^ PDPN^low^ EPCAM^low^ PI^low^ CD31^high^ CD146^high^ cells were sorted as vascular endothelial cells and *Nfat5* knockout was confirmed by qPCR (Supplement 5). Collected cells were resuspended in cryopreservation medium (MLEC medium containing 5% DMSO), stored at -80°C while being shipped to Single Cell Discoveries (Netherlands). Cells were thawed, checked for viability (trypan blue), filtered and processed for scRNAseq analysis (∼2-5,000 cells/sample) according to the indexing protocol CG000315. Sequencing was performed on Illumina Novaseq 6000 (167 GB, 2x50, S1, HMF, up to 50,000 reads/cell). Data analysis BCL files resulting from sequencing were transformed into FASTQ files using 10x Genomics Cell Ranger mkfastq. FASTQ files were mapped using Cell Ranger count. During sequencing, read 1 was assigned 28 base pairs, and was used to identify the Illumina library barcode, cell barcode and UMI. Read 2 was used to map to the mouse reference transcriptome (mm10). Filtering of empty barcodes was done by using Cell Ranger. Data from cells with less than 1000 transcripts/cell or high relative content of mitochondrial genes were removed from further analyses. Data processing for dimensionality reduction (UMAP), cluster identification, differential gene expression and presentation was performed by using BBrowser (V3.5.26, Bioturing, https://doi.org/10.1101/2020.12.11.414136).

### Preparation and analysis of vibratome sections

Mice were euthanized and perfused with 6% gelatine/PBS (37°C) via the right ventricle. Lungs were then inflated by filling the lung with 1% (low melting) agarose/PBS (37°C) via the trachea. After solidification, the lung lobes were removed and fixed in 2% formaldehyde/PBS (4°C) for 18 to 24 hours. Lung lobes were sectioned at 150 µm by vibratome, rinsed in PBS and blocked in PBS containing 5% donkey serum/0.5% Triton X100 for 60 minutes at RT. Subsequently, primary antibodies (goat anti-CD31, AF3628, Biotechne, dilution 1:200; anti-αSMA-Cy3, C6198 Sigma, dilution 1:700; rabbit anti-PDGFB, ab23914, Abcam, Germany) were diluted in PBS containing 5% donkey serum and 0.5% Triton-X100 was added for 24 hours (4°C). After rinsing in PBS, sections were incubated with the secondary antibody (anti-goat-AlexaFluor488, 705-546-147, Dianova, Hamburg, Germany, 1:200) for 24 hours (4°C), rinsed and counterstained with DAPI and mounted with Mowiol. EdU was detected directly by click chemistry according to the manufacturer’s instructions (17-10525, Merck, Darmstadt, Germany). Fluorescence was imaged by using an Olympus IX81 confocal microscope (Olympus) and 3D image stacks were generated from regions of interest containing complete segments of arteries and arterioles. SMC coverage, blood vessel diameter, capillary gap size, and the number of EdU-positive nuclei of at least ten image stacks per lung were assessed using the morphometric tools of the Olympus Xcellence software (version 2.0, build 4768).

### Quantification of PDGF-BB

Supernatants of cultured MLEC were collected and centrifuged at 12,000g for 5 min at 4°C. Supernatants were aliquoted and stored at -80°C until further processing. Lavage fluids from mouse lungs were prepared from mice right after euthanasia by cervical dislocation. A short 23G premade needle catheter was inserted approximately 0.5 cm into the trachea to fill the lungs with 1 ml of Hanks’ balanced salt solution (HBSS) containing 100 µM EDTA and protease inhibitors. The lung lavage solution was then gently aspirated back while massaging the thorax of the mouse. The recovered lung lavage fluid was collected, centrifuged at 12,000 g for 5 minutes at 4°C and stored at -80°C until further processing. PDGF-BB levels in both sample types were measuered (without further dilution) using the Quantikine® Mouse/Rat PDGF-BB Immunoassay ELISA (Cat. #MBB00, Biotechne, Wiesbaden-Nordenstadt, Germany) according to manufacturer’s instructions.

### Determination of the right ventricular systolic pressure (RVSP)

To assess RVSP, mice were anaesthetized with isoflurane (2% conc.) until surgical tolerance was reached (toe reflex test). A pressure-volume catheter (PVR-1030, Millar OEM, Texas, USA) was surgically inserted and advanced through the external jugular vein into the right ventricle. Total blood oxygenation was monitored throughout the procedure with a pulse oximeter (MouseSTAT, Kent Scientific, USA). Pressure pulses were recorded continuously for several minutes (LabChart software, ADI Instruments, USA) once a characteristic stable RV pressure profile was observed. RVSP values were determined by calculating the average of at least ten pressure peaks. Values based on irregular recordings (e.g. technically caused) were excluded from further analysis.

### Echocardiography

Transthoracic echocardiography was performed in anaesthetized mice (1-3% isoflurane and 1 Lpm oxygenated room air) using a Vevo 2100 high-resolution system (Visualsonics, Toronto, ON, Canada) and a 40-MHz MS-550D transducer. Two-dimensional B-mode tracings were recorded in both parasternal long and short-axis views at the level of the papillary muscles and the pulmonary artery (PA), respectively, followed by one-dimensional M-mode tracings in both axes at the level of the papillary muscles or pulsed-wave (PW) Doppler measurement of the peak flow in the PA. The right ventricle (RV) was imaged in B- and M-mode in a modified parasternal long axis view with an adjusted angle focusing on the RV. Data were analyzed offline using VevoLab 3.2.6 and the integrated cardiac measurement package. Ventricular wall thickness at end-diastole was used to characterize RV and LV (left ventricle) anatomy while the change in right- or left-ventricular diameter, respectively, from end-diastole to end-systole was used to assess the contractility and calculate RV or LV fractional shortening (FS). Pulmonary artery hypertension was estimated using the ratio of pulmonary artery acceleration to ejection time in the PW diagram as previously described ^19^. RV cardiac output was monitored using the integral of PA flow, PA diameter and heart rate. M-mode imaging and heart rate were used to calculate LV cardiac output. Three consecutive cardiac cycles were used for each analysis.

### Statistical analysis

Statistical analyses were performed by applying GraphPad Prism (Ver 9.2, GraphPad Software, San Diego, CA, USA). Unless stated otherwise, all results are expressed as mean ± SD. Outliers were identified using the Grubbs’ (α=0.05) or ROUT (Q=1%) test. Statistical evaluation of differences among normally distributed values of two experimental groups was performed using Student’s *t*-test for paired or unpaired data. Statistical evaluation of differences in one parameter between normally distributed values of three or more experimental groups was analyzed by one-way ANOVA with Šídák’s multiple comparisons test or two-way ANOVA with Tukey’s multiple comparisons test, as appropriate. Values of *p*<0.05 were considered statistically significant.

## RESULTS

### Hypoxia stimulates translocation of NFAT5 into the nucleus of murine lung EC

Based on earlier findings ^9^, we assumed that NFAT5 controls transcriptional responses in hypoxia-exposed MLEC. Initially, we investigated NFAT5 activation in hypoxic human umbilical vein endothelial cells (HUVEC), which typically requires its translocation from the cytoplasm into the nucleus. As evidenced by immunofluorescence analyses, low levels of NFAT5 were preferentially located in the cytoplasm under normoxia while NFAT5 was detectable in ∼80% of the EC nuclei 24 hours after exposure to hypoxia (Supplement 6). Corresponding experiments were performed with cultured MLEC, which were exposed to hypoxia for 4, 8 and 24 hours. *Nfat5* expression was not much altered at early time points but significantly increased after 24 hours, which was comparable to the course *Vegfa, a* known HIF1A target (Supplement 7A). Likewise, the protein abundance of NFAT5 as well as its accumulation in the nuclei was increased in hypoxia-exposed MLECs (Supplement 7B-D).

### Loss of Nfat5 in EC affects the transcriptome of the lung after short-term but not long-term exposure to hypoxia

We assumed that lung EC require NFAT5 for adapting their transcriptome to cope with hypoxic stress. To assess the relevance of this process for lung structure and function, we generated a mouse line for EC-specific knockout of *Nfat5*. To this end, we crossed *Cdh5*-CreERT2 transgenic mice with *Nfat5^fl/fl^* mice ^16^ to delete the *Nfat5* gene in EC (*Nfat5^(EC)-/-^*) by tamoxifen treatment followed by a recovery time of at least two weeks. Miglyol (solvent)-treated mice served as controls (*Nfat5^fl/fl^*). Mice were exposed to normoxia or normobaric hypoxia (10% oxygen) for 7 and 21 days. Quantitative PCR analyses indicated a significant increase in *Nfat5* expression in *Nfat5^fl/fl^*but not *Nfat5^(EC)-/-^* mouse lungs (Figure 1A, Supplement 8) and isolated MLEC after exposure to hypoxia for 7 days (Figure 1B). However, *Nfat5* expression was not much altered in lungs exposed to hypoxia for 21 days (Figure 1A). Results from fluorescence-activated cell sorting (FACS) and IF-based analyses suggested that the proportion of MLEC in relation to the total cell population and their structural organization is not altered by genetic ablation of *Nfat5* (Figure 1C and D). Subsequent microarray-based analyses revealed that loss of endothelial *Nfat5* has little impact on the expression pattern of mouse lungs exposed to normoxia or hypoxia for 21 days when comparing *Nfat5^(EC)-/-^* and *Nfat5^fl/fl^* mice as evidenced by the Pearson correlation coefficient (Figure 1E and G). However, knockout of *Nfat5* preferentially altered (early) transcriptional responses to hypoxia after seven days (Figure 1F) correlating with the elevated *Nfat5* expression level at that time point. Gene set enrichment analyses (GSEA) of microarray data from hypoxia (7 d)-exposed *Nfat5^fl/fl^* and *Nfat5^(EC)-/-^* mouse lungs suggested the enrichment of genes associated with ribosomal activity, oxidative phosphorylation and extracellular matrix receptor interaction while those regulating propanoate metabolism, cell cycle and mRNA surveillance were down-regulated (Figure 2H). Additionally, no changes in gene expression associated with smooth muscle contraction (normalized enrichment score (NES): -1.21, adj. p-value: 0.2817) and HIF1α activity (NES: -1.14, adj. p-value: 0.38) were observed in hypoxia (7 d)-exposed *Nfat5^(EC)-/-^*versus *Nfat5^fl/fl^* lungs. The same applies to genes involved in the regulation of immune responses such as cytokines and their receptors (NES: -0.9, adj. p-value: 0.84) or those regulated by NFκB (NES: -1.23, adjusted p-value: 0.29). Most of these gene sets regulated by NFAT5 after 7 days of hypoxia were not altered when comparing *Nfat5^(EC)-/-^* and *Nfat5^fl/fl^* mouse lungs exposed to normoxia or hypoxia for 21 days (Figure 1H). Likewise, no significant changes in the expression of individual genes were observed under these conditions (Supplement 9).

**Figure 1:**
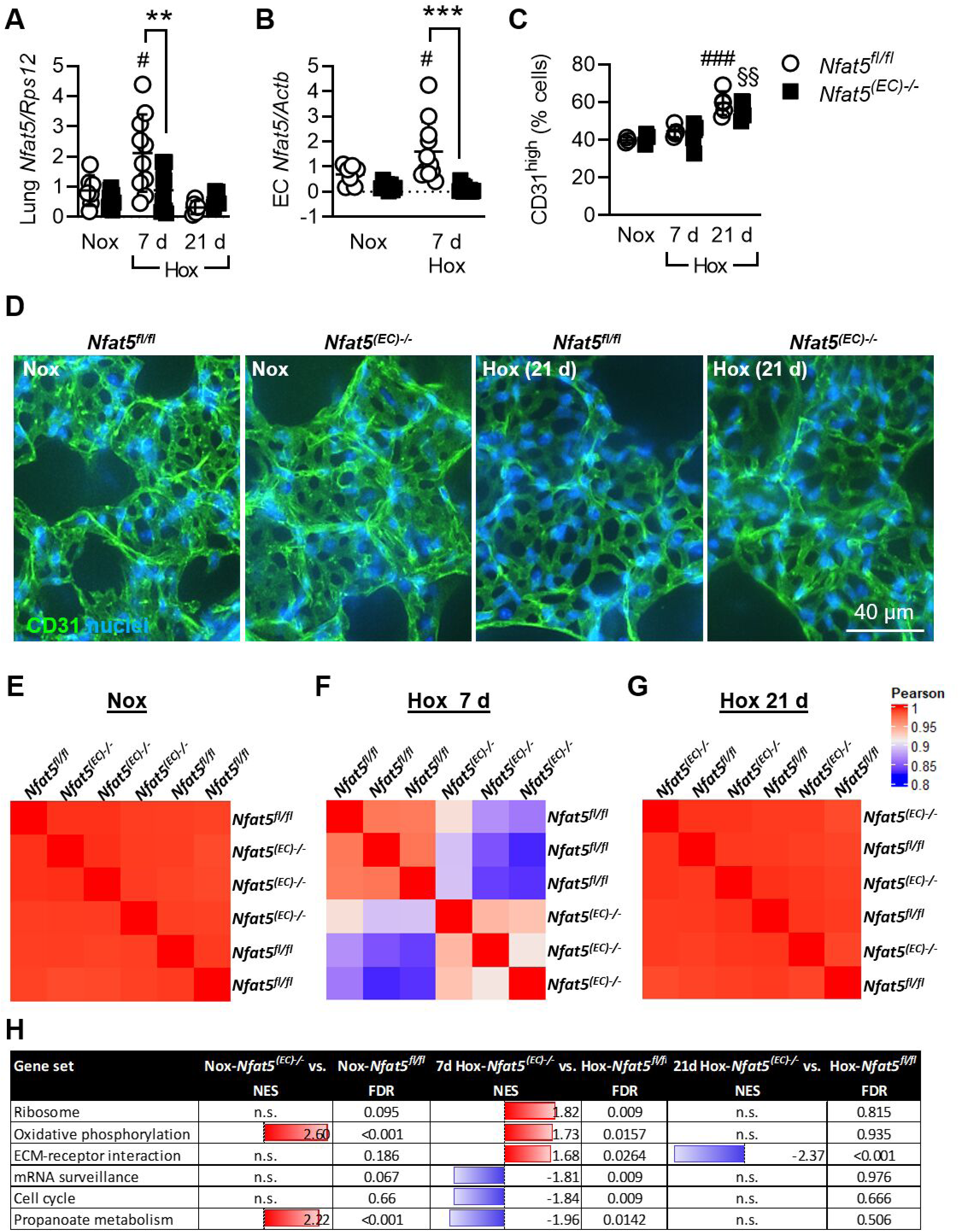
Gene expression profiles of mouse lungs exposed to normoxia/hypoxia for 7 and 21 days upon knockout of *Nfat5* in ECs. (**A**) *Nfat5* RNA expression (qPCR) in whole lungs (#*p*<0.05 vs. Nox-*Nfat5^fl/fl^*, \*\**p*<0.01 as indicated, n=5-10, two-way ANOVA/Tukey’s multiple-comparisons test) and **(B)** EC freshly isolated from lung tissue (^#^*p*<0.05 vs. Nox-*Nfat5^fl/fl^*, \*\**p*<0.01 as indicated, n=7-14, two-way ANOVA/Tukey’s multiple-comparisons test). (**C**) FACS analysis showing the percentage of the EC population in relation to other lung cell populations (^###^*p*<0.001 vs. Nox-*Nfat5^fl/fl^*, ^§§^*p*<0.01 vs. *Nfat5^f(EC)-/-^*, n=3-5, two-way ANOVA/Tukey’s multiple-comparisons test). (**D**) Confocal microscopy/immunofluorescence-based images of the structure of alveolar capillaries (PECAM-1: green, nuclei: blue). (**E-G**) Heatmaps showing the Pearson correlation of gene expression in *Nfat5^(EC)-/-^ versus Nfat5^fl/fl^* lungs after Nox/Hox exposure for 7 and 21 days. (**H**) Enrichment analyses (Kyoto Encyclopedia of Genes and Genomes (KEGG) database, NES - normalized enrichment score, FDR - false discovery rate) showing the top 3 up/down-regulated gene sets in the different experimental groups (Gene sets were selected based on the comparison (7d) Hox-*Nfat5^(EC)-/-^ vs. Hox-Nfat5^fl/fl^*).

**Figure 2:**
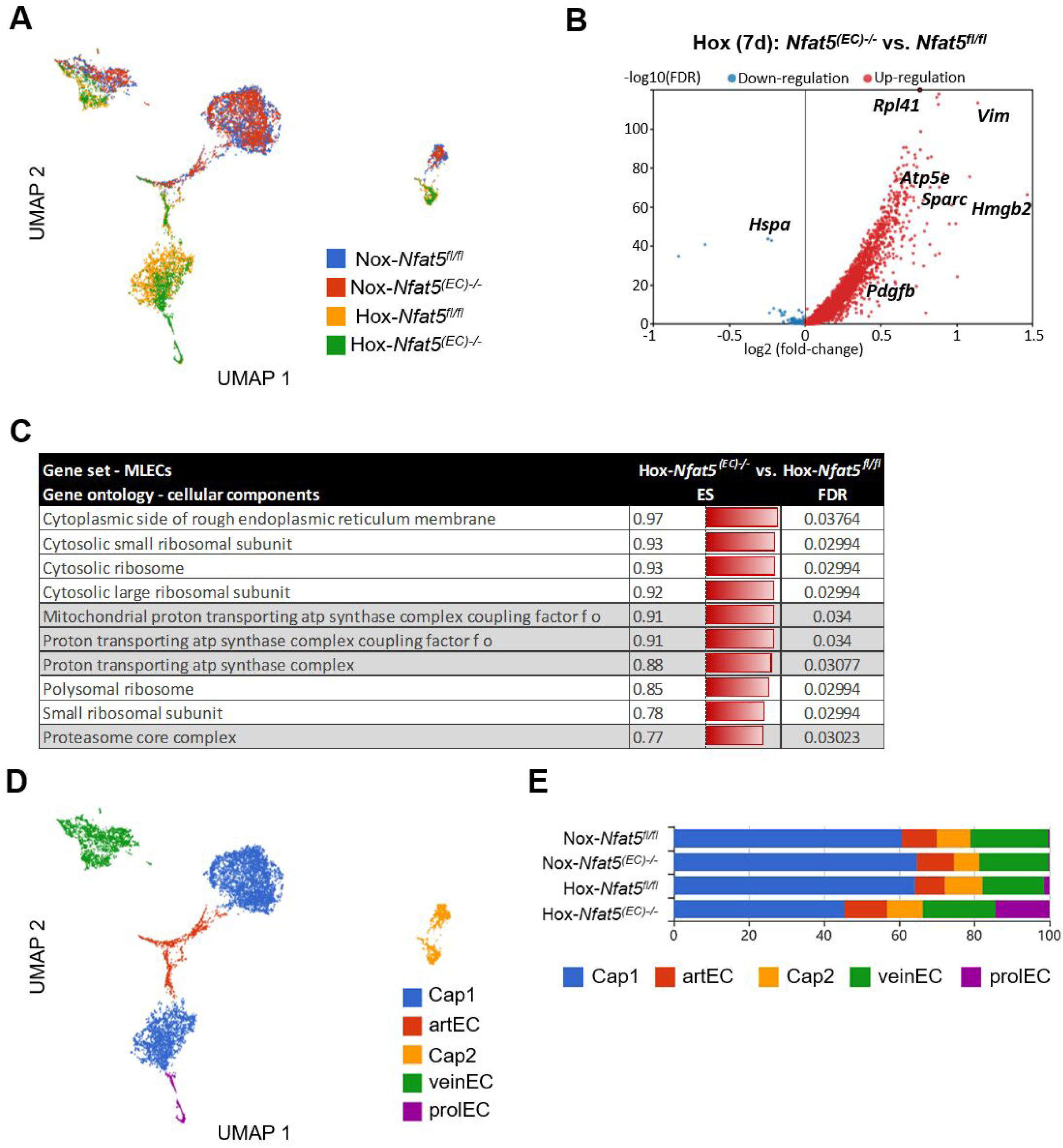
scRNA-seq of MLECs from Nox/Hox (7 d)-exposed *Nfat5^fl/fl^* and *Nfat5^(EC)-/-^ lungs*. (**A**) Uniform manifold approximation and projection (UMAP)-based plot (2 dimensions, 30 components) showing the dimension-reduced expression data of individual cells under four experimental conditions. (**B**) Volcano plot of differentially expressed genes (DEGs) comparing hypoxia (7 d)-exposed *Nfat5^(EC)-/-^ and Nfat5^fl/fl^* MLECs. (**C**) Enrichment analyses (Gene ontology (GO) database, ES - enrichment score, FDR - false discovery rate) showing the top ten regulated gene sets comparing Hox-*Nfat5^(EC)-/-^ and Hox-Nfat5^fl/fl^* MLECs. (**D**) UMAP plot identifying five individual EC clusters (see Supplement 8 for details). (**E**) Relative size (% of total) of each EC subpopulation under different experimental conditions.

In contrast, hypoxia (7 d)-exposed *Nfat5^(EC)-/-^* versus *Nfat5^fl/fl^* mouse lungs showed a decrease in the expression of several endothelial marker genes (e.g. *Cdh5, Pecam1*) although neither the proportion of MLEC, nor the histological organization of the alveolar capillaries was altered under these conditions as described above. Expression of VSMC marker genes (e.g. *Myh11, Acta2*) was not changed. However, *Mir143* - a microRNA that supports VSMC differentiation ^20^ - was down-regulated upon loss of endothelial NFAT5. In line with the GSEA results, elevated levels of transcripts encoding components of the extracellular matrix (e.g. *Col1a*) were detected in *Nfat5^(EC)-/-^* mice after 7 days of hypoxia. Regulation of inflammation-associated genes was marginally altered, which is supported by FACS analyses indicating only minor changes in leukocyte populations of hypoxia (7 d)-exposed lungs from *Nfat5^(EC)-/-^* mice as compared to *Nfat5^fl/fl^*mice (Supplement 10). Interestingly, the expression level of transcripts encoding transforming growth factor beta1 (TGFβ1) and PDGFB - important determinants of tissue remodeling processes - was elevated in hypoxic *Nfat5^(EC)-/-^*versus *Nfat5^fl/fl^* lungs.

### Analysis of gene expression in EC subpopulations by scRNA-seq

The observed changes in gene expression may be attributed to any cell type in the lung. To delineate EC-specific hypoxia (7 d)- and NFAT5-dependent transcriptional responses, we isolated MLEC and processed them for single cell RNA sequencing (scRNA-seq). Uniform manifold approximation and projection (UMAP)-based dimension reduction resulted in a mirror-inverted arrangement of statistically related EC clusters from the normoxia and hypoxia treatment groups (Figure 2A). Comparison of the gene expression of all EC clusters from Nox-*Nfat5^(EC)-/-^* and Nox-*Nfat5^fl/fl^* lungs revealed minor differences (Supplement 11A) supporting the results of the corresponding microarray analysis. Hypoxia significantly altered the transcriptome of *Nfat5^fl/fl^* EC clusters and promoted the expression of many hypoxia-regulated genes such as *Edn1* (endothelin-1)*, Ldha* (lactate dehydrogenase A) and *Pdgfb* (Supplement 11B). Surprisingly, loss of *Nfat5* under hypoxic conditions stimulated gene expression at large as can be deduced from the asymmetric volcano plot of differentially expressed genes (DEGs) of EC clusters in the Hox-*Nfat5^(EC)-/-^* and Hox-*Nfat5^fl/fl^*group (Figure 2B). Subsequent GESA of Hox-*Nfat5^(EC)-/-^* versus Hox-*Nfat5^fl/fl^* EC clusters revealed that gene sets associated with energy and protein metabolism were upregulated (Figure 2C) as was also shown by corresponding analyses of the microarray data.

We next determined marker genes of statistically related EC clusters (Supplement 12), which included recently reported genetic markers of distinct MLEC subpopulations^21–23^. In total, five distinct MLEC subpopulations were annotated including type 1 capillary EC (Cap1), type 2 capillary EC (Cap2), arterial EC (artEC), venous EC (veinEC) and EC with high transcript levels of proliferation markers (prolEC) (Figure 2D). Endothelial cells identified as *Nrp2^high^ Prox1^high^* Mmrn1^high^ lymphatic cell population (49 cells in total) were excluded from further analyses.

Enrichment analyses of gene sets correspondent to those identified in the whole lung indicated that hypoxia has only a minor effect on gene expression associated with cell-matrix interactions, proliferation or RNA surveillance in *Nfat5^fl/fl^* MLEC subpopulations (Supplement 13). Gene expression associated with pyruvate metabolism was up-regulated while genes linked with oxidative phosphorylation were down-regulated. However, the knockout of endothelial *Nfat5* in hypoxia-exposed MLEC stimulated the expression of genes associated with ribosome activity, oxidative phosphorylation, cell-matrix interactions and proliferation in (almost) all subpopulations (as compared to Hox-*Nfat5^fl/fl^*). For the most part these results support the microarray data and indicate that *Nfat5*-deficient MLEC display an elevated level of cellular activation in a hypoxic environment.

With regard to the differential expression of individual genes, each MLEC subpopulation showed a largely specific pattern of top 10 up/down-regulated genes in response to both hypoxia and loss of *Nfat5* (Supplement 14 and 15). For instance, elevated levels of *Edn1* and *Sparcl1* (encodes the matricellular protein hevin) expression was only found in Hox-versus Nox-*Nfat5^fl/fl^* Cap1 and artEC populations. When comparing Hox-*Nfat5^(EC)-/-^* and Hox-*Nfat5^fl/fl^* MLEC, Cap1 and artEC increase the expression of vimentin (*Vim*) - a regulator of angiogenic EC activation ^24^ - while veinEC elevate the expression of endothelial adhesion molecules *Selp*, *Icam1* and *Vcam1*. As a common response of all *Nfat5^fl/fl^* EC subpopulations to hypoxia, we identified *Hspa*-type chaperones, the expression of which was suppressed in hypoxic MLEC after knockout of *Nfat5*.

Considering the results obtained so far, loss of *Nfat5* forced MLECs to particularly express genes suitable for supporting their energy and protein metabolism under hypoxic conditions. Moreover, evaluation of relative changes in the size of MLEC subpopulations implied that the proportion of the Cap1 population was reduced and the prolEC population was enlarged in hypoxia-exposed *Nfat5^(EC)-/-^* versus *Nfat5^fl/fl^* mice (Figure 2E). Subsequent trajectory analyses of the Hox*-Nfat5^(EC)-/-^*EC population identified prolEC as rich in *Sema3c* – a marker of the Cap1 EC population (Supplement 16). Furthermore, expression of selected DEGs (*Atp5e, Cox4i1, Vim, Pdgfb and Cxcl12*) showed strongest correlation with *Sema3c* (Supplement 17A) and GESA indicated increased expression of energy and protein metabolism-associated genes in this subpopulation of MLEC (Supplement 17B). Interestingly, histological analyses of hypoxia (7 d)*-*exposed *Nfat5^(EC)-/-^* and *Nfat5^fl/fl^*mouse lungs did not reveal significant differences in the number of proliferating MLEC (Supplement 18) suggesting that expression of corresponding marker genes may follow compensatory needs.

### Knockout of endothelial Nfat5 stimulates PDGFB release in the hypoxic mouse lung and cultured MLEC

Collectively, our transcriptome analyses implied that *Nfat5* deficiency stimulates metabolism- and proliferation-associated gene expression in MLECs exposed to a hypoxic environment. Furthermore, this appears to be accompanied by remodeling processes as evidenced by the enrichment of genes associated with extracellular matrix components and cell adhesion. We assumed that *Nfat5*-deficient EC may directly or indirectly drive hypoxia-mediated remodeling processes by paracrine mediators such as TGFβ1 and PDGFB, which were up-regulated in hypoxic *Nfat5^(EC)-/-^*versus *Nfat5^fl/fl^* mouse lungs. While *Tgfb1* expression showed a similar upregulation, that was, however, restricted to a small proportion of hypoxic *Nfat5^-/-^* (versus Hox-*Nfat5^fl/fl^*) Cap1 EC, *Pdgfb* expression was evident in the bulk of the Cap1 population and significantly increased in Hox-*Nfat5^-/-^* versus Hox-*Nfat5^fl/fl^*Cap1 ECs (Figure 3A). In contrast, hypoxic *Nfat5*-deficient artEC slightly down-regulated *Pdgfb* expression (as compared to Hox-*Nfat5^fl/fl^*). To validate these findings, RNA from FAC-sorted MLEC and lung tissue was analyzed by qPCR. *Pdgfb* but not *Tgfb1* was up-regulated in hypoxic *Nfat5^-/-^* (versus *Nfat5^fl/fl^*) MLEC (Figure 3B-D). Corresponding analyses of hypoxic lung tissue confirmed that knockout of endothelial *Nfat5* stimulated the expression of *Pdgfb* (Figure 3E and F). Likewise, immunofluorescence-based detection techniques revealed elevated PDGFB levels in alveolar capillaries of hypoxia-exposed *Nfat5^(EC)-/-^* versus *Nfat5^fl/fl^* mouse lungs (Supplement 19), which was supported by analyses of bronchoalveolar lavage fluids (Figure 3G). In addition to *Tgfb1* and *Pdgfb*, expression of the artEC population marker *Cxcl12* ^23^ was upregulated in hypoxic *Nfat5^-/-^* (versus *Nfat5^fl/fl^*) Cap1 EC (Supplement 20A). However, neither microarray-based results nor qPCR analyses of FAC-sorted lung EC indicated a significant impact on the overall *Cxcl12* expression level at the investigated time point (Supplement 20B).

**Figure 3:**
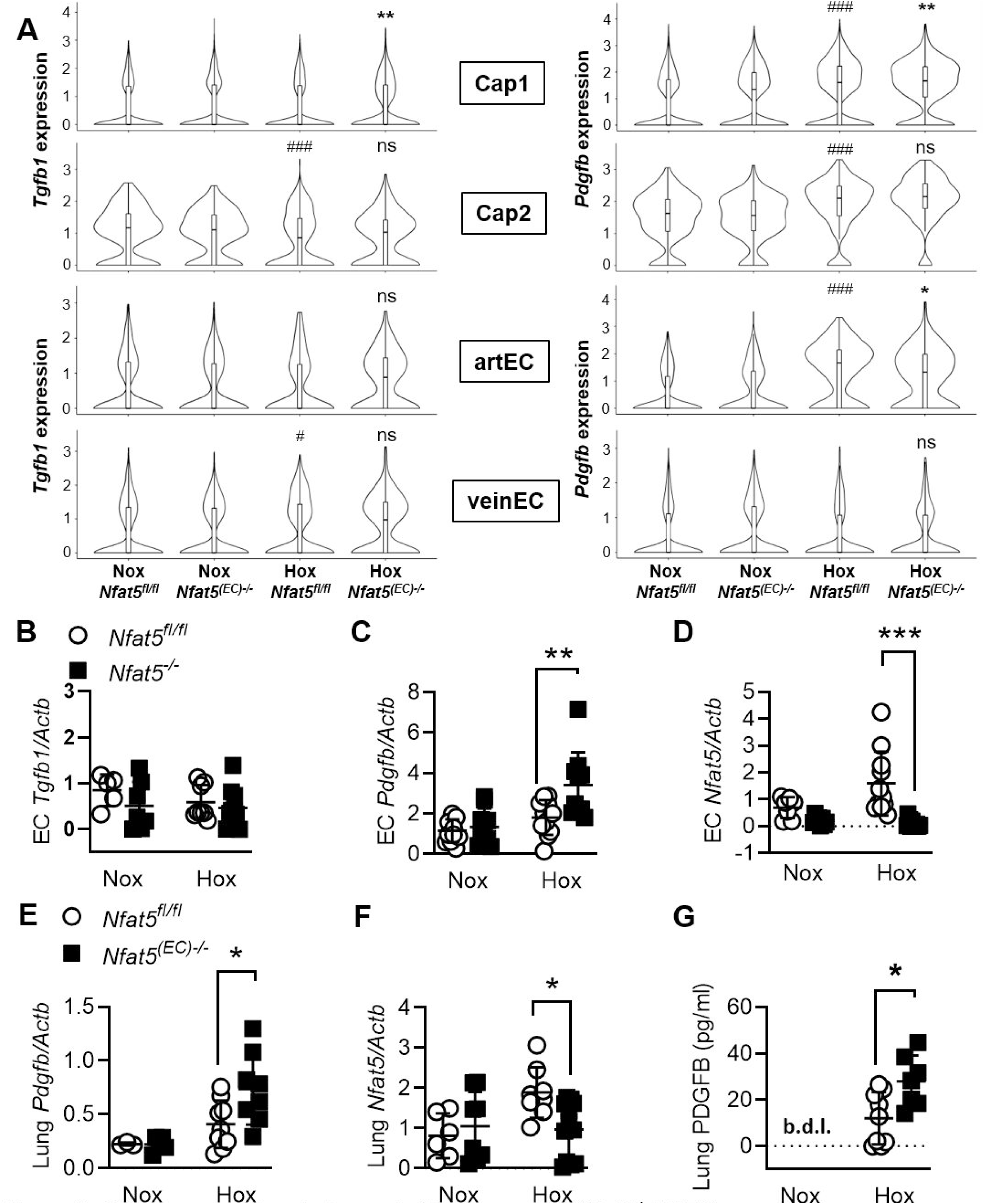
*Pdgfb* expression in hypoxia (7 d)-exposed *Nfat5^-/-^* MLECs. (**A**) Violin plots showing the condition-dependent distribution of the *Pdgfb* expression data in EC subpopulations (bandwith calculation by Silverman’s rule of thumb, boxes show interquartile range, \**p*<0.05, \*\**p*<0.01 and ns (no significance, *p*>0.05,) vs. Hox-*Nfat5^fl/fl^*; ^###^*p*<0.001 vs. Nox-*Nfat5^fl/fl^*, one-way ANOVA/Tukey’s multiple-comparisons test). (**B-F**) qPCR analysis of *Tgfb1*, *Pdgfb* and *Nfat5* expression in (FAC-sorted) MLECs (B-D) and whole lung tissue (E, F) after exposure to normoxia or hypoxia for 7 d (\**p*<0.05, \*\**p*<0.01, \*\*\**p*<0.001 as indicated, n=3-14, two-way ANOVA/Tukey’s multiple-comparisons test). (**G**) ELISA of PDGFB levels in lung lavage fluids (b.d.l. - below detection limits) from both *Nfat5^M^f* and *Nfat5E^C^-* mice after exposure to normoxia or hypoxia for 7 d (\**p*<0.01 as indicated, n=4-9, two-way ANOVA/Tukey’s multiple-comparisons test).

We next cultured MLEC isolated from *Nfat5^fl/fl^* mouse lungs to investigate the mechanism that modifies *Pdgfb* expression. The cell culture model mimicked the *in vivo* findings as exposure to hypoxia amplified *Pdgfb* expression and the release of PDGFB in *Nfat5^-/-^*versus *Nfat5^fl/fl^* MLEC (Figure 4A-C). Although stimulation of gene expression upon knockout of a transcription factor seems counterintuitive, NFAT5 has been reported to alter transcriptional activity by modulating DNA methylation and chromatin accessibility ^25^ as well as by inhibiting recruitment of transcription factors to enhancer regions of gene promoters ^26^. As our results did not substantiate an impact of NFAT5-mediated DNA methylation (Supplement 21) or chromatin accessibility (Supplement 22) on *Pdgfb* expression under the tested experimental conditions, we investigated whether binding of HIF1α - a transcription factor that is crucial for hypoxia-induced *Pdgfb* expression in MLEC (Supplement 23) ^6^ - to the *Pdgfb* promoter is altered in the presence or absence of NFAT5. By *in silico* analysis of the corresponding promoter region (Figure 4D), we identified a putative NFAT5 binding site (NP-4) that overlaps with a HIF1α binding site (HRE - hypoxia response element). We assessed the binding of both HIF1α and NFAT5 to these regions by applying chromatin immunoprecipitation. Interaction of NFAT5 with NP-4 was increased after exposure of MLECs to hypoxia for 24 hours and abolished in *Nfat5*-deficient MLECs. HIF1α showed an enhanced association of with this promoter region in absence of NFAT5 (Figure 4E). NFAT5 may thus simply interfere with the binding of HIF1α to (at least) one of its binding sites in the *Pdgfb* promoter region. In turn, over-expression of *NFAT5* diminished the expression of *Pdgfb* in hypoxia-exposed *Nfat5^fl/fl^*MLEC (Supplement 24) suggesting that raising the abundance of NFAT5 is sufficient to suppress *Pdgfb* expression.

**Figure 4:**
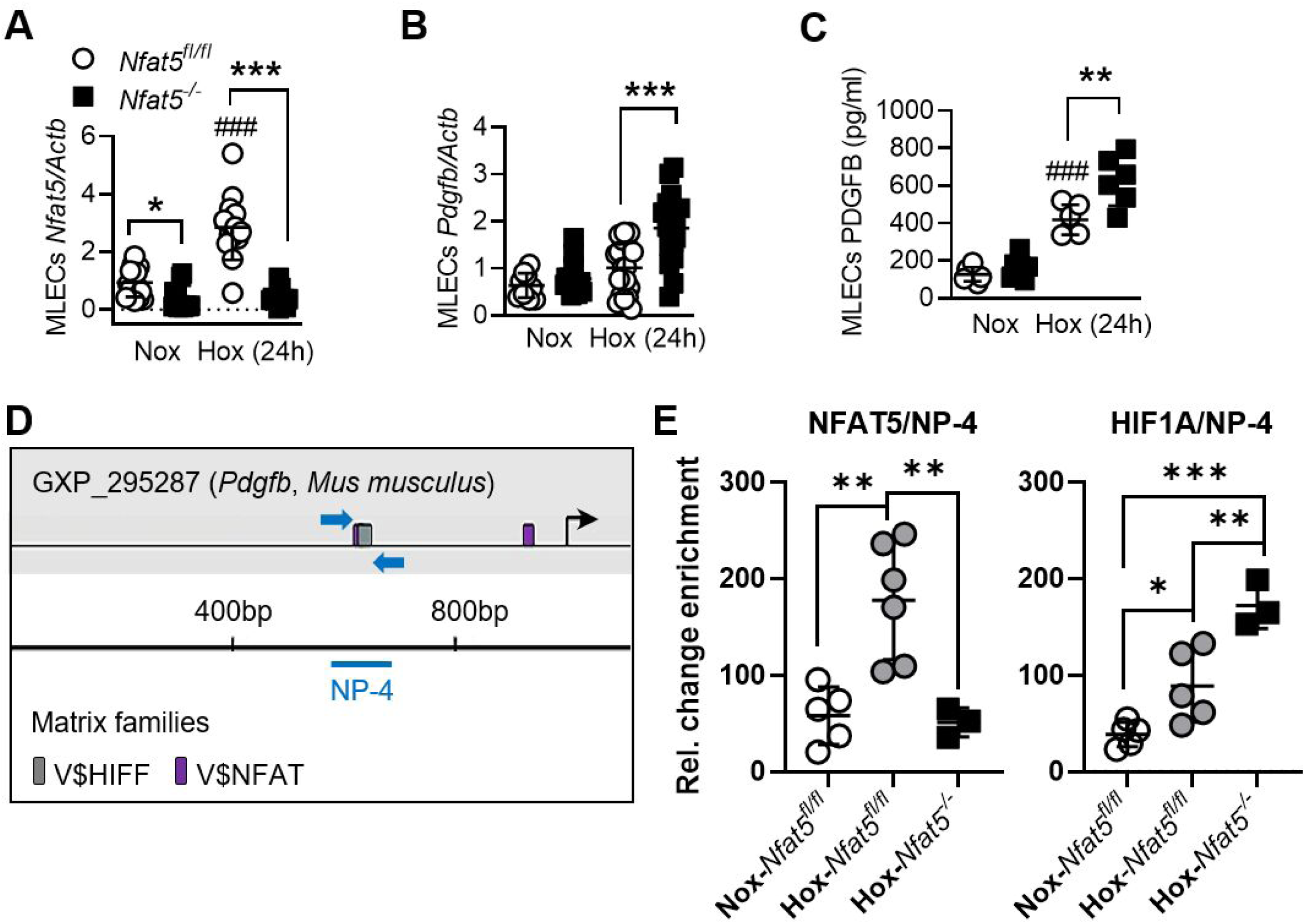
NFAT5-modulated HIF1a binding to the *Pdgfb* promoter region. (**A-C**) qPCR analyses of *Nfat5* and *Pdgfb* expression (A, B) and ELISA-based assessment of PDGFB levels in supernatants (**C**) of *Nfat5^ff/ff^* and *Nfat5^-/-^* in MLECs exposed to normoxia and hypoxia for 24h (^###^*p*<0.001 vs. Nox-*Nfat5^ff/ff^*, \*\**p*<0.01, \*\*\**p*<0.001 as indicated, n=3-9, two-way ANOVA/Tukey’s multiple-comparisons test). (**D**) Illustration of binding sites for NFAT5 (V$NFAT) and HIF1a (V$HIFF) in the promoter region of *Pdgfb* as calculated by Genomatix (software suite v3.13, MatInspector, Precigen Bioinformatics, Munich, Germany). Binding of transcription factors was tested for a sequence located in region NP-4 (blue). (**E**) Chromatin isolated from *Nfat5^ff/ff^* and *Nfat5^-/-^* normoxia- and hypoxia-exposed MLECs was precipitated by anti-NFAT5 and anti-Hifla antibodies and subjected to ChlP-qPCR analysis (ns - no significance (*p*>0.05), \**p*<0.05, \*\**p*<0.01, \*\*\**p*<0.001, as indicated, one-way ANOVA/Tukey’s multiple-comparisons test, n=3-6).

### Nfat5 deficiency in EC elevates right ventricular systolic pressure (RVSP) and coverage of pulmonary arterioles by VSMCs

Elevated PDGFB levels may promote proliferation of VSMCs in the hypoxic lung ^6^ and subsequently enhance their coverage of small arterioles. In fact, morphometric analyses of vibratome sections of mouse lungs by confocal microscopy revealed an increase in the proliferation of smooth muscle actin (αSMA)-positive cells in *Nfat5^(EC)-/-^*versus *Nfat5^fl/fl^* hypoxic (7 d) mouse lungs (Figure 5A). Considering the characteristics of the applied mouse model, such remodeling processes become usually functionally overt within three weeks after onset of hypoxia. Moreover, the degree of muscularization of distal arterioles was shown to directly increase the resistance of pulmonary arteries at this time point ^6^. We assessed effects of endothelial NFAT5 on pulmonary hemodynamics by determining the right ventricular systolic pressure (RVSP) in *Nfat5^(EC)-/-^* and *Nfat5^fl/fl^*mice exposed to normoxia and hypoxia for 7 and 21 days. RVSP was increased in all hypoxia-exposed animals, whereby highest values were observed in *Nfat5^(EC)-/-^* mice after 21 days (Figure 5B). Morphometric IF-based analyses showed augmented VSMC coverage of small lung arterioles in *Nfat5^(EC)-/-^* versus *Nfat5^fl/fl^* mice after being exposed to hypoxia for 21 days (Figures 5C). In line with the microarray data indicating the up-regulation of gene expression associated with the extracellular matrix in Hox-*Nfat5^(EC)-/-^*mouse lungs, we also observed a pronounced perivascular deposition of collagen – suggesting a higher degree of (peri)vascular remodeling in hypoxia-exposed *Nfat5^(EC)-/-^* versus *Nfat5^fl/fl^*lungs (Supplement 25).

**Figure 5:**
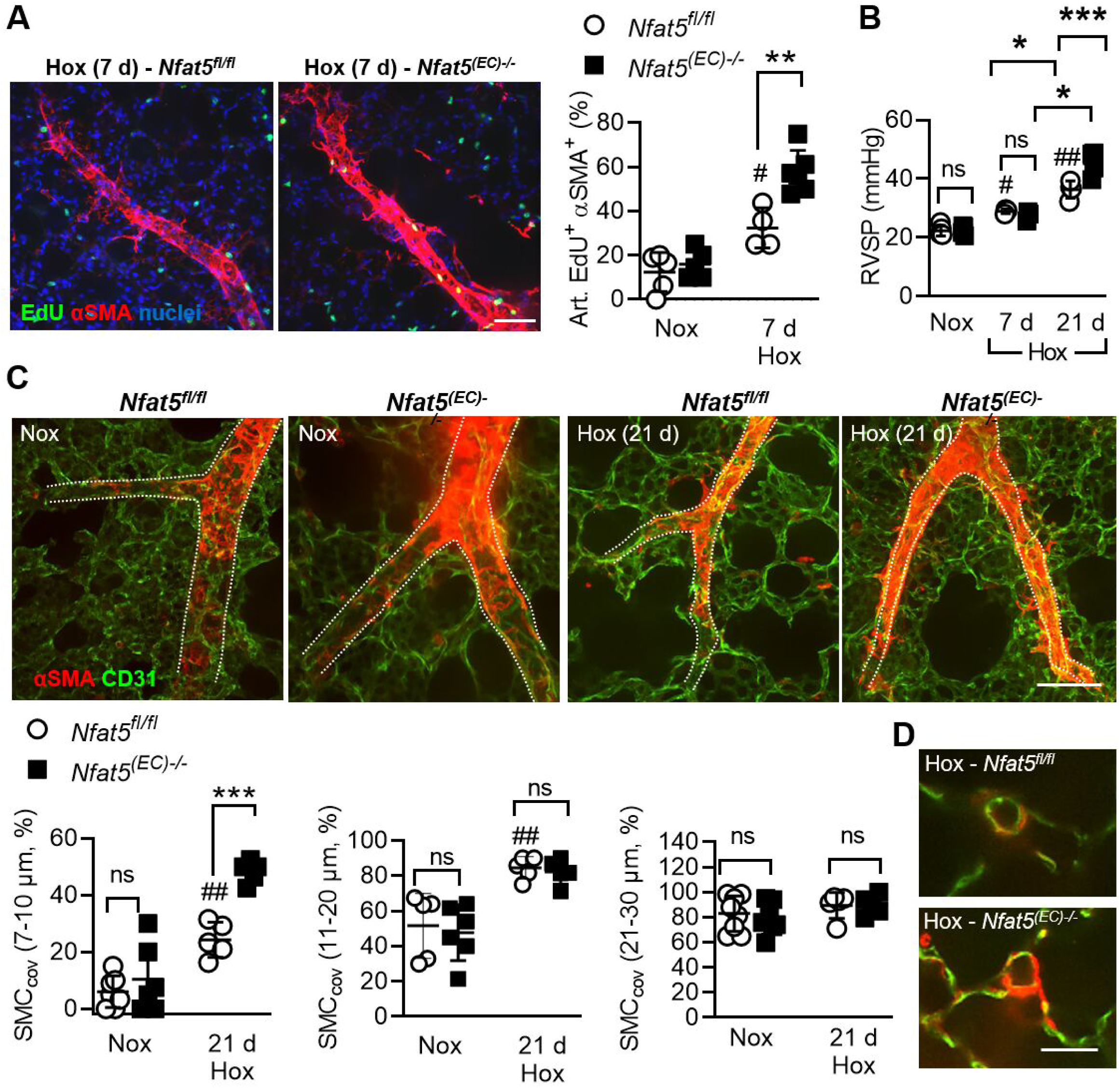
Analysis of VSMC proliferation and coverage of pulmonary arterioles in hypoxia-exposed *Nfat5^(EC)-/-^* mice. (**A**) Confocal microscopy/IF-based detection of the proliferation marker EdU (dotted lines indicate the route of the arteries/arterioles, aSMA: red; Edu: green, nuclei: blue, scale bar: 50 pm) in vibratome sections of mouse lungs from both *Nfat5f^f/^f^f^* and *Nfat5^(EC)-/-^* mice. The percentage of EdU^+^ aSMA^+^ cells located in small arteries (diameter ∼20-50 pm) was calculated (n=4-5). (**B**) Assessment of the right ventricle systolic pressure (RVSP) in normoxia- and hypoxia-exposed *Nfat5^ff/ff^* and *Nfat5^(EC)-/-^* mice (n=3-5). (**C**) Confocal microscopy/immunofluorescence-based analysis of VSMC coverage of arterioles/arteries (SMC_cov_, percentage of aSMA artery area) with different diameter ranges (aSMA, red; CD31: green, scale bar: 50 pm, n=4-5). ^#^*p*<0.05, ^##^*p*<0.05 vs. Nox-*Nfat5^fl/fl^*; \**p*<0.05, \*\**p*<0.01, \*\*\**p*<0.001, ns – no significance (*p*>0.05) as indicated, two-way ANOVA/Tukey’s multiple-comparisons test). (**D**) Representative images of cross-sections of small arterioles comparing the level of VSMC coverage between *Nfat5^fl/fl^* and *Nfat5^(EC)-/-^* arterioles (scale bar: 20 pm).

### Loss of endothelial Nfat5 impairs right ventricular functions in hypoxia-exposed mice

Chronically elevated RVSP values may negatively affect cardiac functions. Echocardiographic analyses of normoxia-exposed *Nfat5^(EC)-/-^* and *Nfat5^fl/fl^* mice showed neither structural nor functional (Figure 6A-H) abnormalities. Likewise, no changes in the left ventricular (LV) or right ventricular (RV) cardiac output or LV wall thickness (Figure 6A-D) were observed after exposure of the animals to hypoxia for 21 days. However, both the thickness of the intraventricular septum (IVS) and the wall of the right ventricle were increased in Hox-*Nfat5^(EC)-/-^* as compared to Hox-*Nfat5^fl/fl^* mice (Figure 6E and F). In addition, we observed a decrease in RV fractional shortening (correlating with RV contractility) and a pronounced reduction in the pulmonary artery acceleration/ejection time ratio (inversely related to pulmonary vascular resistance) in hypoxia-exposed *Nfat5^(EC)-/-^*versus *Nfat5^fl/fl^* mice (Figures 6G and H). While the functional data of hypoxia-exposed *Nfat5^fl/fl^* mice reflect typical characteristics of this model ^27, 28^, loss of endothelial NFAT5 aggravated hypoxia-induced pulmonary hypertension as well as the impairment of (RV) cardiac functions, presumably at least in part due to increased distal artery muscularization. Hence, in a hypoxic environment adequate transcriptional and functional responses of lung EC critically rely on NFAT5 to restrict their capacity for initiating detrimental vascular remodeling processes.

**Figure 6:**
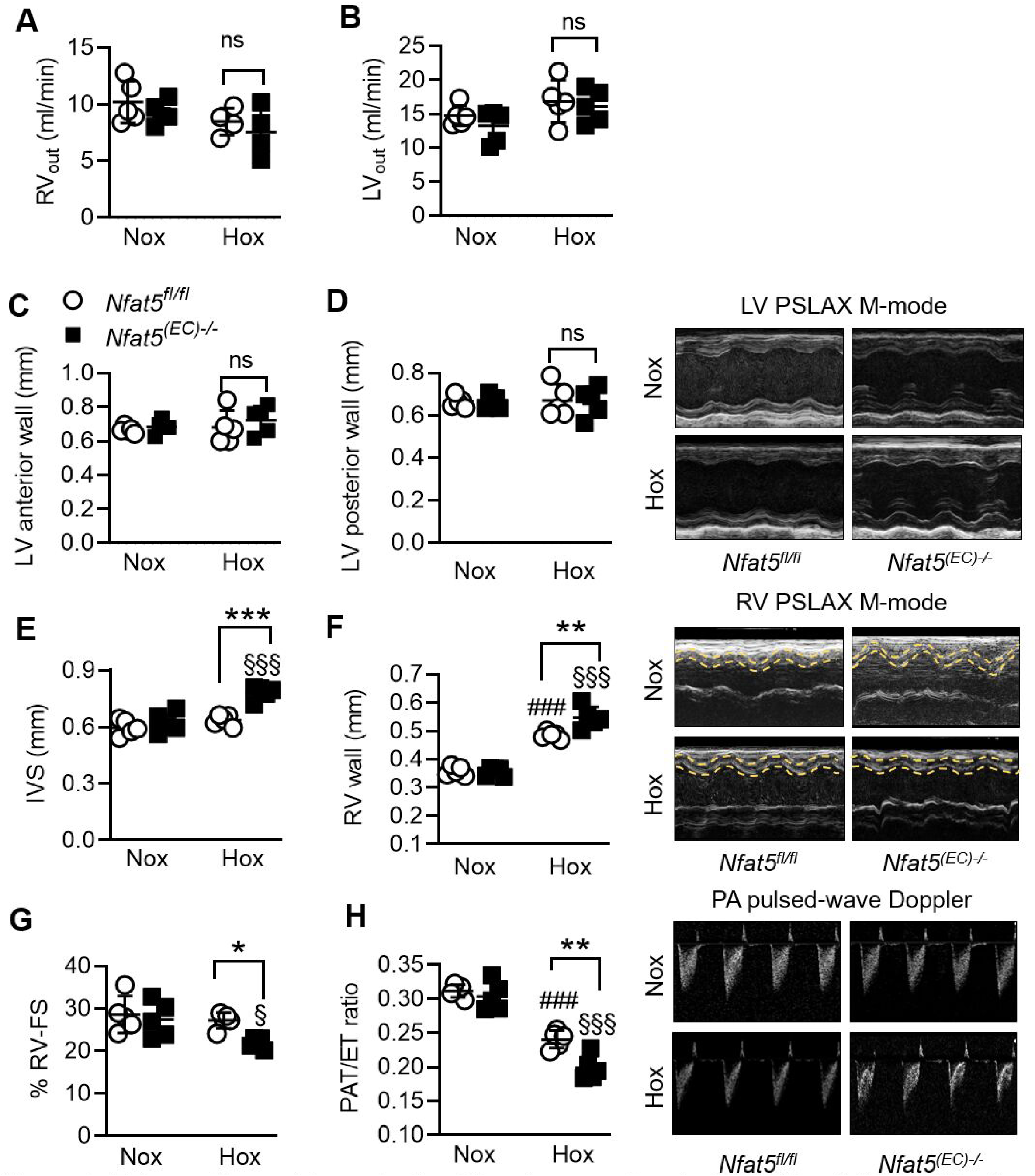
Echocardiographic analysis of heart parameters in *Nfat5^fl/fl^* and *Nfat5^(EC)-/-^* mice exposed to normoxia/hypoxia for 21 days. (**A**) Right ventricular (RV) and (**B**) left ventricular (LV) cardiac output (CO) (the difference between RVCO and LVCO is due to methodology), (**C**) LV anterior wall thickness, (**D**) LV posterior wall thickness (representative images show the LV parasternal long axis view (PSLAX) in motion mode), (**E**) interventricular septum (IVS) thickness, (**F**) RV wall thickness (representative images show the RV PSLAX in motion mode), (**G**) RV fractional shortening indicating the RV systolic function, (**H**) pulmonary artery acceleration time (PAT) to ejection time (ET) ratio (representative images show Doppler-based pule waves of the pulmonary artery (PA)). \**p*<0.05, \*\**p*<0.01, \*\*\**p*<0.001 as indicated; ns - no significance (*p*>0.05) as indicated, ^#^*p*<0.05, ^##^*p*<0.01, ^###^*p*<0.001, vs. Nox-*Nfat5^fl/fl^* ; ^§^*p*<0.05, ^§§^*p*<0.01, ^§§§^*p*<0.001, vs. Nox-*Nfat5^(EC)-/-^*, two-way ANOVA/Tukey’s multiple-comparisons test, n=4-5.

## DISCUSSION

Obstruction of the respiratory tract or hypoventilation cause a local or global drop in ventilation of the lungs. As a consequence, the partial pressure of oxygen decreases in the affected areas including both the alveolar and vascular space. Such a scenario creates a hypoxic environment and develops in patients suffering from COPD or ventilation disorders. In the long run, hypoxia drives structural remodeling processes in the lung, such as the coverage of distal arterioles by VSMCs. This progressively increases pulmonary vascular resistance, and eventually leads to the development of PAH and impairs right ventricular function. While the long-term effects of chronic hypoxia are well documented, short-term responses of MLEC to hypoxia are less well characterized. However, their relevance should not be underestimated as mouse model-based studies suggest that mechanisms deciding about histological and functional alterations observed three weeks after the onset of hypoxia (structural remodeling, RVSP, cardiac functions), are already initiated during the first week^7^. Our findings suggest that MLEC adapt their transcriptome to cope with the hypoxic environment within this period thereby exhibiting common as well as subpopulation-specific responses. Based on marker gene expression, we distinguished between four different main clusters of EC, which correspond to those already described in the literature^21–23^. Besides markers attributed to arterial and venous EC, the expression pattern of Cap1 EC resembles that of ‘general’ capillaries, which may act as progenitor EC in homeostasis and repair ^22^. The transcript profile of Cap2 EC identify them as ‘aerocytes’ reported to support gas exchange and trafficking of leukocytes. After being exposed to hypoxia for seven days, all MLEC up-regulated pyruvate synthesis-associated genes but down-regulated genes associated with oxidative phosphorylation, reflecting a typical adaptation of the cellular energy metabolism to low oxygen levels. Proliferation and adhesion-associated genes were preferentially expressed by artEC while prostanoid-related gene expression was particularly diminished in hypoxic versus normoxic venous EC. Interestingly, enrichment of genes associated with leukocyte tethering and rolling was detected in all but Cap2 EC, which, however, did not much alter the composition of the leukocyte population in the lung at the investigated time point. With respect to specific genes, all MLEC increased the expression of *Hspa1a* and/or *Hspa1b* in response to hypoxia underlining the general relevance of heat shock protein 70 (HSP70) for protein homeostasis under stress-related conditions ^29^. Interestingly, both Cap1 and artEC exclusively elevated the expression level of *Sparcl1* encoding the matricellular SPARC-like protein 1 that is supposed to induce endothelial quiescence and limit angiogenesis ^30^. This finding confirms data from recent scRNA-seq analyses of MLEC at later stages of hypoxia showing that *Sparcl1* is preferentially expressed in the capillary and lung arterial endothelium ^23^.

From a pathophysiological point of view, hypoxia acts as an important environmental stressor forcing MLEC to rapidly adapt their phenotype. Considering the impact of NFAT5 on the expression pattern of hypertonicity-exposed ^31, 32^ or biomechanically stimulated cells ^33, 34^, we assumed that while it is not critically involved in maintaining cellular homeostasis, it may contribute to stress-related transcriptional changes in response to hypoxia. In fact, the conditional endothelium-specific knockout of *Nfat5* in mice exposed to normoxia evoked no obvious impairment of cardiovascular functions supporting earlier data obtained from mice with smooth muscle cell-specific *Nfat5* deficiency ^9, 35^. In contrast, global knockout of *Nfat5* in mice causes renal atrophy and early lethality ^36^ demonstrating its pivotal role in regulating adaptive responses in cells exposed to osmotic stress.

Hypoxia stimulated *Nfat5* expression and nuclear translocation of the protein in EC as observed in VSMCs after being exposed to biomechanical^33^, metabolic^37^ or hypoxic stress^9^. Entry of NFAT5 into the nucleus requires multiple posttranslational modifications such as its phosphorylation by c-Abl (ABL1) and/or JNK in osmotically or biomechanically stressed cells^31, 38, 33, 34^. Although the exact hypoxia-induced molecular modification of NFAT5 has not yet been investigated, it is likely that JNK contributes to its activation in hypoxic lung EC^39, 40^. Moreover, palmitoylation of proteins - another modification of NFAT5 that may support its nuclear translocation ^34^ - was demonstrated in hypoxia-exposed pulmonary arteries ^41^.

The fundamental relevance of NFAT5 for adjusting the MLEC transcriptome may be deduced from the consequences of its knockout. Notably, lack of *Nfat5* affected gene expression after short-term (7 d) but not much after long-term (21 d) exposure to hypoxia, which was also reflected by both its expression level and the degree of changes in the lung transcriptome at these time points. Although the transcriptional activity of NFAT5 in hypoxia-exposed MLEC appears to be temporary, its deficiency had nevertheless deleterious consequences on structure and function of the lung and heart. The most likely cause for the increase in RVSP and impaired RV structure and function is the extensive coverage of distal pulmonary arteries by VSMC, resulting in a corresponding increase in pulmonary vascular resistance.

In general, knockout of endothelial *Nfat5* robustly promoted or extended the expression of multiple genes in hypoxia-exposed MLEC. For instance, ribosome and oxidative phosphorylation-associated gene sets were enriched in Hox-*Nfat5^(EC)-/-^* versus Hox-*Nfat5^fl/fl^* lungs as substantiated by both microarray and scRNAseq-based analyses suggesting an elevated or extended demand of *Nfat5*-deficient MLEC for energy-consuming protein turnover. Similar observations were made in *Nfat5*-deficient pulmonary artery VSMCs, which boosted oxidative phosphorylation-associated gene expression and mitochondrial respiration in mice exposed to hypoxia for 7 days ^9^. In the Cap1 EC population this appeared to be accompanied by a substantial increase in expression of proliferation markers.

Counterintuitively, loss of NFAT5 had a general multifaceted stimulatory impact on gene expression. Paradoxically, *in silico* analyses suggested that most genes up-regulated in hypoxia-exposed *Nfat5*-deficient MLEC exhibit no corresponding binding sites in their promoters. Accordingly, a substantial part of the transcriptomic alterations observed in *Nfat5^-/-^*MLEC may originate from indirect compensatory responses to the impaired expression of rate-limiting genetic determinants required to adjust the cellular phenotype.

Besides the general effects on gene regulation, we specifically analyzed the impact of NFAT5 on the expression of *Pdgfb* due to its critical role in the remodeling of the pulmonary vasculature. In this context, the applied mouse model strongly relies on the release of PDGFB from the lung endothelium to stimulate migration and proliferation ^6^ of PDGF receptor β-expressing VSMCs, which was shown to peak seven days after the onset of hypoxia ^7^. Our results herein suggest that both arterial and capillary EC contribute to the PDGFB level in the hypoxic lung. While we may not fully exclude other cell types such as macrophages as PDGFB source^42^, additional scRNA-seq analyses of CD45+ lung cell populations did not indicate a significant contribution of alveolar macrophages relating thereto at the investigated time point (data not shown). Likewise, the population size of these immune cells was not altered in Hox-*Nfat5^(EC)-/-^* versus Hox-*Nfat5^fl/fl^* lungs suggesting - together with the transcriptome data - that *Nfat5* regulates no major immunomodulatory functions in MLEC exposed to hypoxia for seven days despite its fundamental capacity to enhance the transcriptional activity of NFκB in LPS-stimulated cells ^43^.

Strikingly, knockdown of *Nfat5* increased expression and release of PDGFB in MLEC and/or lungs of Hox-*Nfat5^(EC)-/-^*versus Hox-*Nfat5^fl/fl^* mice providing a mechanistic explanation for the elevated level of proliferating pulmonary VSMCs observed after seven days of hypoxia and their increased coverage of distal arterioles after 21 days. Although *Cxcl12* expression may also contribute to the activation of pulmonary VSMCs ^44^ and was shown to be significantly stimulated in Hox-*Nfat5^(EC)-/-^*versus Hox-*Nfat5^fl/fl^* MLEC, the confirmation of the expression data was not fully convincing. This is in line with observations indicating a comparable expression level of *Cxcl12* in capillary and arterial MLEC under normoxic and hypoxic conditions. In contrast, the rise in *Pdgfb* expression in hypoxia-exposed *Nfat5^-/-^* versus *Nfat5^fl/fl^ MLEC* was confirmed *in vivo* and *in vitro*. Hypoxia-associated transcription of this gene in EC is to the most part dependent on HIF1α ^45^. Correspondingly, interference with HIF1α expression decreases *Pdgfb* expression in MLEC, limits VSMC coverage of distal pulmonary arterioles and consequently lowers RVSP in hypoxia-exposed mice ^6^. Here, inhibition of HIF1α completely abolished expression of *Pdgfb* in cultured hypoxia-exposed MLEC (Supplement 22). Considering the relevance of HIF1α for *Pdgfb* expression, we assumed that loss of NFAT5 indirectly restricted its global transcriptional activity. However, neither the protein level of HIF1α was altered in hypoxia-exposed *Nfat5^-/-^* MLEC, nor the expression of several other HIF1α targets such as *Vegfa*.

NFAT5 may basically affect gene expression by i) direct stimulation ^46^, ii) indirect enhancement ^25, 43^ and iii) suppression ^47, 48^ including epigenetic mechanisms involving NFAT5-dependent DNA methylation and modification of chromatin accessibility. However, none of them appeared to substantially affect *Pdgfb* expression in the investigated context as suggested by our results. We therefore assumed that NFAT5 may directly influence HIF1α- controlled transcription of *Pdgfb* as supposed for other transcriptions factors acting as cooperation or co-activation partners ^49^. In line with this view, binding of NFAT5 to one binding site located in the *Pdgfb* promoter region displaced HIF1α from an adjacent hypoxia response element (HRE). Although we cannot exclude other possible effects directly or indirectly evoked by *Nfat5* deficiency as modulators of *Pdgfb* expression in hypoxic MLEC, we consider direct interference of NFAT5 with the transcriptional activity of HIF1α as the most plausible explanation.

In summary, our findings suggest that NFAT5 is indispensable for the adequate and rapid adjustment of transcriptional responses of MLEC exposed to low oxygen levels. Consequently, loss of *Nfat5* demands additional and extended efforts from the endothelium to cope with a hypoxic environment forcing the expression of genes, which preferentially promote metabolic activity, growth and *Pdgfb* expression of capillary MLEC. This dysfunctional adaptation process exacerbated VSMC coverage of distal pulmonary arterioles with detrimental impact on the right ventricular afterload and heart function. Our findings delineated so far unknown context- and cell-dependent regulatory features of NFAT5, which are crucial for the compensation of hypoxia-mediated stress. Moreover, they underline the relevance of temporary adaptive processes of the lung endothelium for the outcome and progression of chronic hypoxia-related diseases.

## Data availability

The data underlying this article are available in the article and in its Supplementary material online. The raw microarray data underlying this article are available in the Gene Expression Omnibus (GEO) database as record GSE230055 at https://www.ncbi.nlm.nih.gov/geo/query/acc.cgi?acc=GSE230055, and can be accessed with the following token for review: evanwmcwrzwfxmv

While under review, the scRNA-seq data underlying this article will be shared on reasonable request to the corresponding author. They will be made available to the public in the course of the publication process.

## Acknowledgements

The authors thank for the excellent support by Maria Harlacher, Paula Breuer and Caroline Arnold. We thank Single Cell Discoveries for their help with project design, single-cell sequencing services and data analysis.

## Non-standard Abbreviations and Acronyms

αSMA: smooth muscle actin
ArtEC: arterial endothelial cell
ATAC-seq: transposase-Accessible Chromatin using sequencing
Cap1: type 1 capillary endothelial cell
Cap2: type 2 capillary endothelial cell
CMV: cytomegalovirus
COPD: chronic obstructive pulmonary disease
DEGs: differentially expressed genes
EC: endothelial cells
Edn1: endothelin-1
EdU: 5-ethynyl-2′-deoxyuridine
FACS: fluorescence-activated cell sorting
FCS: fetal calf serum
FS: fractional shortening
GSEA: gene set enrichment analyses
HIF1A: hypoxia-inducible factor 1 alpha
HPV: pulmonary vasoconstriction
HRE: hypoxia response element
HUVEC: human umbilical vein endothelial cells
KLF4: krüppel-like (transcription) factor 4
Ldha: lactate dehydrogenase A
LV: left ventricle
MACS: magnetic activated cell sorting
MLEC: murine lung endothelial cells
Nfat5: nuclear factor of activated T-cells 5
PAH: pulmonary arterial hypertension
Pdgfb: platelet-derived growth factor B
RV: right ventricle
RVSP: right ventricular systolic pressure
Sparcl1: SPARC-like protein 1
TGFβ1: transforming growth factor beta1
TonEBP: tonicity enhancer binding protein
VeinEC: venous endothelial cell
Vim: vimentin
VSMC: vascular smooth muscle cell

